# Mutation of the Galectin-3 Glycan Binding Domain (*Lgals3*-R200S) Enhances Cortical Bone Expansion in Male and Trabecular Bone Mass in Female Mice

**DOI:** 10.1101/2020.01.09.900787

**Authors:** Kevin A. Maupin, Daniel Dick, VARI Vivarium, Transgenics Core, Bart O. Williams

## Abstract

The study of galectin-3 is complicated by its ability to function both intracellularly and extracellularly. While the mechanism of galectin-3 secretion is unclear, studies have shown that the mutation of a highly conserved arginine to a serine in human galectin-3 (*LGALS3*-R186S) blocks glycan binding and secretion. To gain insight into the roles of extracellular and intracellular functions of galectin-3, we generated mice with the equivalent mutation (*Lgals3*-R200S) using CRISPR/Cas9-directed homologous recombination. Consistent with a reduction in galectin-3 secretion, we observed significantly reduced galectin-3 protein levels in the plasma of heterozygous and homozygous mutant mice. We observed a similar increased bone mass phenotype in *Lgals3*-R200S mutant mice at 36 weeks as we previously observed in *Lgals3*-KO mice with slight variation. Like *Lgals3*-KO mice, *Lgals3*-R200S females, but not males, had significantly increased trabecular bone mass. However, only male *Lgals3*-R200S mice showed increased cortical bone expansion, which we had previously observed in both male and female *Lgals3*-KO mice and only in female mice using a separate *Lgals3* null allele (*Lgals3*). These results suggest that the trabecular bone phenotype of *Lgals3*-KO mice was driven primarily by loss of extracellular galectin-3. However, the cortical bone phenotype of *Lgals3*-KO mice may have also been influenced by loss of intracellular galectin-3. Future analyses of these mice will aid in identifying the cellular and molecular mechanisms that contribute to the *Lgals3*-deficient bone phenotype as well as aid in distinguishing the extracellular vs. intracellular roles of galectin-3 in various signaling pathways.

## Introduction

Galectin-3 (*Lgals3*) is a chaperone protein which functions in numerous cell processes both intracellularly via protein-protein interactions and extracellularly by binding specific glycans on glycoproteins [1]. The extracellular interactions are important for polarized glycoprotein secretion [2], as well as glycoprotein turnover [3]. Intracellularly, galectin-3 supports subcellular localization of various binding partners [4]. The function of galectin-3 in all of these compartments hampers our ability to identify functional mechanisms based upon galectin-3 knockdown or overexpression studies. For example, galectin-3 regulates β-catenin protein levels in multiple cell lines [5-18], however it is not clear whether this is through direct intracellular interaction between galectin-3 and β-catenin [8, 15] or indirectly potentially through an extracellular binding partner [11, 16, 19, 20]. It is not even clear whether galectin-3 interacts directly with β-catenin, as some papers have stated an inability to co-IP galectin-3 with β-catenin [7, 9] and another paper has shown co-IP of extracellular galectin-3 with β-catenin if a membrane impermeable chemical crosslinking agent was used [20]. In order to make progress in addressing galectin-3 mechanisms, we need to be able to limit the function of galectin-3 in one or more cellular compartments.

There is a great interest in targeting galectin-3 in several pathologies, particularly fibrosis [21-27]. We previously showed that genomic deletion of galectin-3 in mice (*Lgals3*-KO) protected them against bone loss due to aging [28] or sex-hormone deprivation[29]. Therefore, galectin-3 inhibitors might also be useful for preventing osteoporosis. Better understanding of the intracellular vs. extracellular functions of galectin-3 could lead to smarter drug development. For example, it could prove more efficacious to target extracellular galectin-3, while leaving intracellular galectin-3 function unperturbed. Generating an animal model with a mutant version of galectin-3 that preferentially disrupts extracellular galectin-3, would greatly aid in addressing this hypothesis.

Studies show that it is possible to mutate galectins to selectively disrupt their glycan-dependent functions, while potentially preserving their intracellular functions. Mutation of a highly conserved arginine to a serine in the glycan binding domain of human galectin-3 (R186S) blocks its glycan binding and secretion [30]. While this mutant has thus far only been characterized for its lack of function extracellularly [30, 31], mutation of the structurally equivalent arginine in galectin-7 blocks glycan binding, but still allows galectin-7 to function intracellularly [32, 33]. It is also possible to dissociate galectin-1 glycan binding from intracellular interactions and regulation of mRNA splicing [34]. For these reasons, we chose to create the equivalent R186S mutation in mouse galectin-3 (*Lgals3*-R200S) in order to gain insight into the role of extracellular galectin-3 for the regulation of bone mass.

We were able to successfully generate the *Lgals3*-R200S allele using CRISPR/Cas9 and a single-stranded DNA oligonucleotide as a template for homologous recombination. Like *Lgals3*-KO mice [35], *Lgals3*-R200S mice were grossly normal. Heterozygous mice were fertile and produced wild-type, heterozygous, and homozygous mutant offspring at normal Mendelian ratios. Because we had previously observed significantly enhanced bone mass at 36 weeks in *Lgals3*-KO mice, we also assessed bone mass in *Lgals3*-R200S mice. Like *Lgals3*-KO mice, there was a sex-dependent increase in trabecular bone mass of female *Lgals3*-R200S mice. While we previously observed increased cortical bone expansion in both male and female *Lgals3*-KO mice, only male *Lgals3*-R200S had this phenotype. Similar to *Lgals3*-KO mice, the increased cortical bone expansion was coupled with reduced tissue quality (reduced max stress). However, while we previously observed a strong decrease in tissue level stiffness (elastic modulus) in *Lgals3*-KO mice, *Lgals3*-R200S mice showed no change in tissue or whole bone stiffness values. We believe that the similarities between *Lgals3*-R200S and *Lgals3*-KO mice (i.e., increased trabecular bone mass in female mice and increased cortical bone expansion in males) likely reflect the role of extracellular galectin-3 loss for increasing bone mass. Conversely, we believe that the differences between the two models reflect the role of intracellular galectin-3 (tissue stiffness and lack of female cortical bone expansion). Further studies need to be performed on these mice and their cells to characterize if the mutation affects intracellular functions of galectin-3. However, we believe this new model will be a valuable tool for studying galectin-3 and discriminating the phenotypic effects of the loss of extracellular vs. total galectin-3 function.

## Materials and Methods

### gRNA Design and Synthesis

The sequence encompassing 40 bp upstream and 40 bp downstream from the target mutation site (*Lgals3* codon 200 in exon 5) was probed for all potential sgRNA sites using the Zhang Lab’s Optimized CRISPR Design Tool (crispr.mit.edu). A potential sgRNA binding site with a protospacer adjacent motif (PAM; AGG) was identified 8 bp downstream from codon 200. This sgRNA was selected for its close proximity to the desired mutation site and low predicated probability for off-target binding (quality score 65).

A 20 bp DNA fragment (synthesized by Integrated DNA Technologies (IDT), San Diego, CA) representing the sgRNA plus <u>BbsI</u> complementary overhangs (sgRNA FWD: 5’–<u>CACC</u>TTTGCCACTCTCAAAGGGGA-3’ and sgRNA REV: 5’-<u>AAAC</u>TCCCCTTTGAGAGTGGCAAA-3’) was cloned into a BbsI digested pX330-U6-Chimeric_BB-CBh-hSpCas9 plasmid (42230; Addgene, Cambridge MA). Template DNA for in vitro transcription was generated by PCR amplification of the gRNA sequence—which included both the guide sequence as well as the scaffold sequence encoded in the pX330-U6-Chimeric_BB-CBh-hSpCas9 plasmid—using Phusion HF DNA polymerase and a primer set (synthesized by IDT, San Diego, CA) consisting of a FWD primer recognizing the cloned sgRNA sequence with a <u>T7 RNA polymerase recognition sequence</u> fused on the 5’ end (5’-<u>AATACGACTCACTATAGGG</u>TTTGCCACTCTCAAAGGGGA-3’) and a REV primer that recognized the terminal end of the gRNA scaffold sequence on the pX300 plasmid (5’-AAAAGCACCGACTCGGTGCC-3’). After verifying that the reaction consistently only produced a single product by agarose gel electrophoresis, the template was purified from the reaction mixture using the QIAquick PCR purification kit (Qiagen, Germantown, MD) and eluted with RNAse free water.

In vitro transcription was performed on the purified gRNA template using a MEGAshortscript Kit (Ambion; ThermoFisher Scientific, Waltham, MA) in four 20 μl reactions. The synthesized RNAs were pooled, treated with DNase to remove remaining template, and then purified using a MEGAclear Kit (Ambion; ThermoFisher Scientific, Waltham, MA). Purity of eluted gRNA product was determined by formalin gel electrophoresis and gRNA was quantified on a Nanodrop 2000c (ThermoFisher Scientific, Waltham, MA). Following quantification, gRNA was stored at −80 °C.

### Single Strand Oligonucleotide Donor Template

The single stranded oligonucleotide (ssODN) that was used as a template for homologous recombination (200 bases, PAGE purified) was synthesized by IDT (San Diego, CA). The sequence was: 5’-ATGATGTTGCCTTCCACTTTAACCCCCGCTTCAATGAGAACAACAGGAGAGTCATTGTGTGTAACACGAAGCAGGACAATAACTGGGGAAAGG AAGA<u>GTC</u>ACA<u>A</u>TC<u>T</u>GC<u>G</u>TTCCCCTTTGAGAGTGGCAAACCATTCAAAGTAAGTTGG GGCTTTGGCTGTATGCGCACAGCGTTCTCTTACCAAGGGGAATCACGGAAA-3’. Targeted mismatches are underlined in the ssODN sequence.

### Microinjection and Animal Maintenance

Mouse zygotes were obtained by mating super ovulated B6C3F1/J females with B6C3F1/J males (Jackson Labs, Bar Harbor, ME). RNAs and ssODN were thawed and mixed just prior to injections for final concentrations of: 30 ng/μL WT Cas9 mRNA (Sigma Aldrich, St. Louis, MO), 15 ng/μL gRNA, and 50 ng/μL ssODN. The mix was microinjected into the pro-nuclei of zygotes, and the injected embryos were incubated at 37 °C until they were transferred into pseudo-pregnant females at the two-cell stage. At the end of the injection day, any remaining mix was run out on a formalin gel to confirm that the RNA was not degraded. The mice were housed in climate-controlled conditions (25 °C, 55% humidity, and 12 h of light alternating with 12 h of darkness) and fed a standard LabDiet Rodent Chow 5010 (Purina Mills, Gray Summit, MO). All animal procedures were in compliance with the protocol approved by the Institutional Animal Care and Use Committee at VARI.

### Genotyping

Genomic DNA was isolated from tail clips at weaning and necropsy by proteinase K digestion and ethanol precipitation. Genotypes were assigned by allele specific PCR. Primers were synthesized by Integrated DNA Technologies, San Diego, CA. The PCR reactions included a hot start at 94 °C for 1.5 min followed by 32 amplification cycles of 94 °C for 30 sec, 55 °C for 30 sec, 68 °C for 1.5 min + 3 sec/cycle. These cycles were followed by 68 °C for 5 min. Then the amplified PCR products were analyzed by 2% agarose gel electrophoresis. For the *Lgals3*-R200S reaction, wild-type mice (*Lgals3* +/+) yielded a single 483 bp band, homozygous mutant mice (*Lgals3-*R200S^KIKI^) yielded a 131 bp band and a 586 bp band, and heterozygotes (*Lgals3-*R200S^KI+^) had thee bands of 131 bp, 483 bp, and 586 bp. Sanger sequencing was performed in order to verify that the correction mutations and no spurious indels were present in the *Lgals3-*R200S allele. Tail tip DNA from *Lgals3-*R200S positive animals was first amplified by Phusion HF DNA polymerase PCR using primers encompassing exon 5 of *Lgals3* (FWD: 5’-TTCAGGAGAGGGAATGATGTTG-3’ and REV: 5’-CTGAAGGAGCTGAAGGACAC-3’). The product of this reaction was purified using a QIAquick PCR Purification Kit (Qiagen, Germantown, MD), then cloned into pMiniT Vectors with the NEB PCR Cloning Kit, and transformed into NEB 10-beta Competent E.coli (NEB, Ipswich, MA). Colonies positive for the *Lgals3-*R200S allele by PCR were grown overnight at 37 °C with nutation at 200 rpm. DNA from small aliquots of these cultures were then amplified using the TempliPhi DNA amplification kit and submitted to Genewiz (South Plainfield, NJ) for Sanger sequencing using a forward primer upstream of the insertion site on the pMiniT Vector that came with the NEB PCR Cloning Kit. (5’-ACC TGCCAACCAAAGCGAGAAC-3’).

### Dual Emission X-Ray Absorptiometry (DEXA)

Mice were anesthetized via inhalation of 2% isoflurane (TW Medical Veterinary Supply) with oxygen (1.0 L·min^−1^), weighed, and then placed on a specimen tray in a PIXImus II bone densitometer (GE Lunar) for analysis. Bone mineral density (BMD) and body fat percentage were calculated by the PIXImus software based on the bone and tissue areas in the subcranial region within the total body image.

### Plasma Isolation and Enzyme-Linked Immunosorbent Assays (ELISA)

Immediately following euthanasia, approximately 0.5 mL of whole blood was collected by heart puncture and transferred to a microcentrifuge tube containing 5 μL of 0.5 M Ethylenediaminetetraacetic acid (EDTA) pH 8.0. To separate plasma, tubes were centrifuged at 6,000 xg for 6 min. Plasma galectin-3 from male and female wild-type and heterozygous mice was measured using a DuoSet ELISA Development System kit (D1197; R&D Systems, Minneapolis, MN). Wells were coated with 100 μL of a 2 μg/mL dilution of the galectin-3 capture antibody and only 1 μL of plasma was required per well. The kit included a recombinant galectin-3 that was used for generating a dilution curve for quantification.

### Micro-computed Tomography (μCT)

After euthanasia, right lower limbs and spines were defleshed and fixed in 10% neutral buffered formalin (NBF) for 72 h, rinsed with sterile distilled water, and stored in 70% ethanol at 4 °C. Whole femurs and L3 vertebrae were imaged using a desktop SkyScan 1172 microCT imaging system (SkyScan, Kontich, Germany). Scans for this study were acquired at 80 kV using a 5.98 μm voxel size. The femoral trabecular volume of interest in the aging study encompassed regions from 0.25-1.75 mm from the distal growth plate. For analyses of trabeculae within the body of L3 vertebrae, a 1.5 mm volume centered on the midpoint was used. Cortical measurements were obtained from a 0.6 mm segment that was 45% of the distance proximal of the length of the diaphysis from the growth plate (diaphysis length = distance of femoral head-distance of growth plate). Tissue mineral density and bone mineral density values were obtained using a standard regression line generated by converting the attenuation coefficients to mineral density from scans of hydroxyapatite standards with known densities (0.25 and 0.75 g/cm^3^).

### Mechanical Testing

Following euthanasia, left femurs were defleshed, wrapped in PBS pH 7.2 soaked gauze, and stored at −20 °C. In order to calculate tissue level mechanical parameters, these femurs were thawed at room temperature for 30 min in PBS pH 7.2 and analyzed by μCT at 80 kV with a 9.98 μm voxel size—analyses were performed as described for cortical bone measurements for the right femurs—and then the femurs were returned to −20 °C storage. For mechanical testing, the femurs were thawed a second time and allowed to equilibrate to room temperature for two h in PBS pH 7.2. Following equilibration, a standard four-point bend testing procedure was performed using a TestResources 570L axial-torsional screw-driven testing system (TestResources, Shakopee, MN) with displacement rate of 0.005 mm/s. The distances between the lower and upper supports were 7.3 and 3.5 mm, respectively. The supports had radii of curvature of 0.5 mm at each point of contact with the femur. Displacement was applied by the upper supports in the anterior-posterior direction such that the anterior of the femur was in compression and the posterior was in tension. Force and displacement were directly measured from the load cell and crosshead, respectively. Tissue level mechanical parameters (max stress and elastic modulus) were calculated as described for four-point bending [36] where max stress = (max force*a*c)/(2I_min_) and elastic modulus = stiffness*(a2/(12I_min_))*(3L-4a); where a = the distance between an upper and lower support beam, L is the distance between the lower support beams, I_min_ is the minimum calculated value of inertia, and c is the radius of the bone. I_min_ and c were obtained by μCT.

## Statistical Analyses

A χ^2^ test (df = 5; α = 0.05) was used to verify Mendelian distribution of pups born from *Lgals3*-R200S heterozygous crosses. For most comparisons, the differences between *Lgals3*-wild-type (+/+) and *Lgals3*-R200S (KI/+ and KI/KI) mice were determined using two-way ANOVAs within age groups (sex, genotype). The Holm-Sidak method was used in post hoc analyses to identify significant differences (α = 0.05). However, due to the large difference in plasma Gal3 levels between male and female mice, one-way ANOVAs with Dunnett’s post hoc tests were performed within sex between the genotypes.

## Results

### Generation of *Lgals3-*R200S Allele using CRISPR/Cas9

Mutation of a conserved arginine to a serine in the glycan binding groove of human galectin-3 (*LGALS3-*R186S) prevents galectin-3 secretion and glycan binding [30, 31, 37]. The generation of a mouse with retained intracellular galectin-3 that is expressed normally, but does not function extracellularly, would greatly aid in understanding the role of galectin-3 in each cellular compartment. To generate such a mouse, we mutated the homologous arginine to serine in the mouse *Lgals3* gene using CRISPR/Cas9 (*Lgals3*-R200S; Figure 1A).

**Figure 1.**
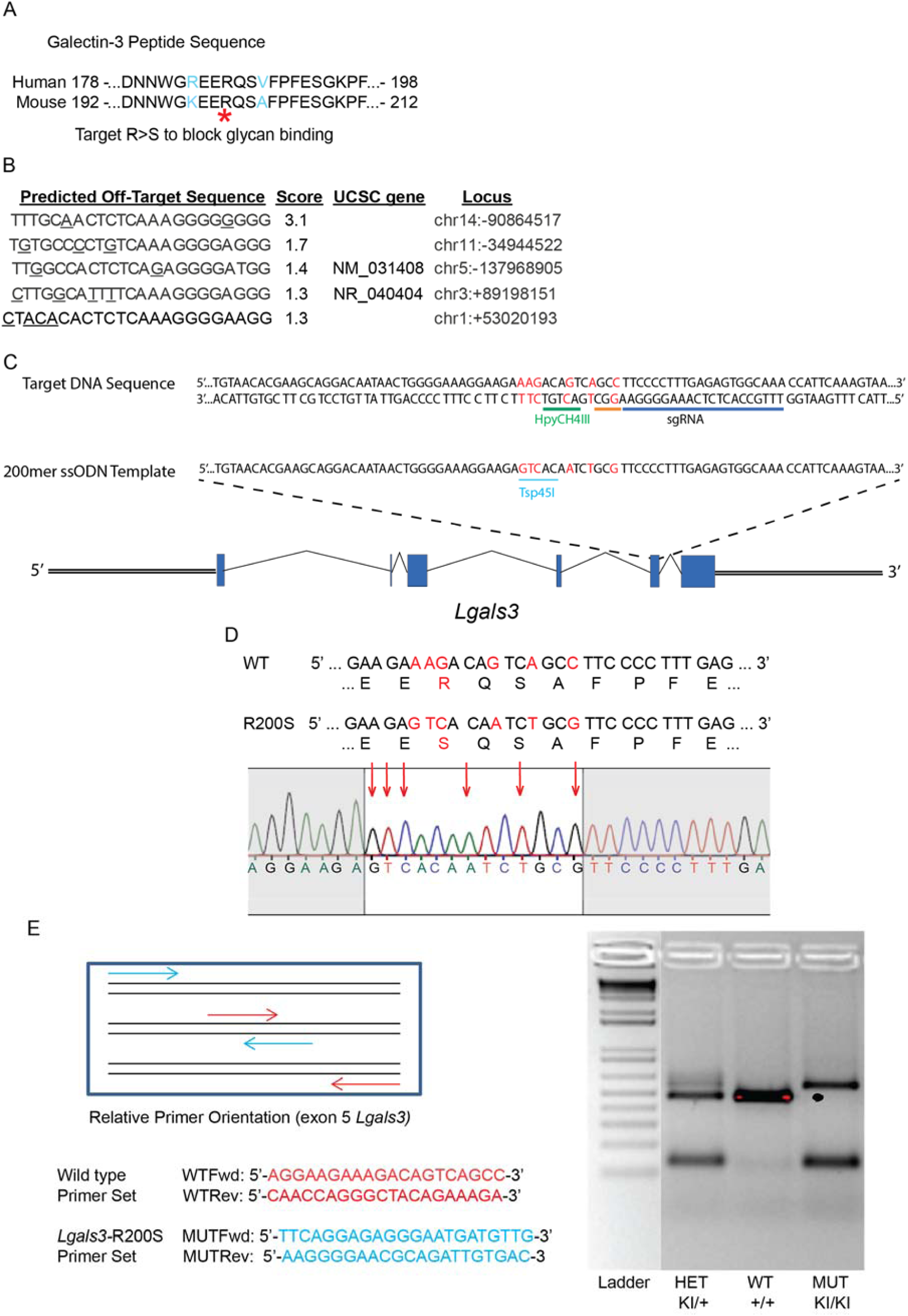
Generation of *Lgals3*-R200S knock-in allele by CRISPR/Cas9. (A) Comparison of human and mouse galectin-3 amino acid sequence and identification of arginine to mutate to generate the glycan binding deficient galectin-3. (B) Predicted off-target binding sites for single guide RNA (sgRNA). (C) Target sequence of *Lgals3* gene showing where sgRNA would bind. Red letters indicate nucleotides to be mutated following homologous recombination of the single stranded oligonucleotide (ssODN) template. (D) Sanger sequencing confirmation of incorporation of mutations. (E) PCR mediated identification of mice with wild type and *Lgals3*-R200S alleles. WT = wild type; MUT = mutant.

To accomplish our targeted mutation, we first identified the closest potential gRNA binding site using the Zhang Lab’s Optimized CRISPR Design Tool (crispr.mit.edu). This gRNA binding site had low predicted off-target binding sites (Figure 1B) with a protospacer adjacent motif (PAM) that was within 8 bp of codon 200 in *Lgals3.* In order to introduce our mutation via homologous recombination, we designed a 200 bp single stranded oligo donor (ssODN) template by substituting the arginine codon (AGA) with a serine (TCA). In order to make identification of the *Lgals3-*R200S allele easier by PCR, we included additional silent mutations. These silent mutations also disrupted a unique HpyCH4III restriction site and generated a novel Tsp45I restriction site which could be used for additional genotyping strategies (Figure 1C). We identified 3 out of 120 pups that were heterozygous for our targeted mutation as identified by allele specific PCR of tail clip DNA from weanlings. These mice (1 female and 2 males) were mated with wild-type C57BL/6J mice to determine if the mutation could be passed through the germline. The female mouse produced a single litter of 3 pups after several months of failed matings, all of which were wild-type. Due to the low number of pups, we were unable to conclude whether the mutant allele was present in her germ cells. One of the males robustly produced only wild-type pups, suggesting that the *Lgals3-*R200S allele was not incorporated into his germ cells. Thankfully, the second male produced offspring with ∼50% of the pups showing positive PCR results for the *Lgals3-*R200S allele, suggesting that the mutant allele was fully present in his germline. Sanger sequencing revealed that these pups did have the desired mutant allele (Figure 1D). Heterozygous offspring from this F1 generation were crossed to generate the mice analyzed for skeletal phenotypes in this study. PCR genotyping was used to identify wild-type (*Lgals3*^+/+^), heterozygous (*Lgals3*-R200S^KI/+^), and homozygous (*Lgals3*-R200S^KI/KI^) mutant mice (Figure 1E). We continue to backcross the *Lgals3-*R200S allele onto the C57BL/6J background for several generations, to remove any potential off-target modifications for future studies.

### No Major Changes in Body Composition in Aged *Lgals3*-R200S Mice

A total of 179 pups from 21 litters (∼8.5 pups per litter) were weaned and genotyped for this study. Of the 179 pups, 48.6% (87) were male and 51.4% (92) were female. Within males, 28.7% (25) were wild-types, 49.4% (43) were heterozygotes, and 21.8% (19) were homozygous mutants. For females, 19.6% (18) were wild-types, 59.8% (55) were heterozygotes, and 20.7% (19) were homozygous mutants. The observed ratios were not statistically different from expected Mendelian ratios as determined by a χ^2^ test. Analysis of body composition changes by dual-energy X-ray absorptiometry (DEXA) revealed no significant differences for male or female *Lgals3-*R200S mice in weight, area bone mineral density (aBMD), or bodyfat percentage (%BF) as shown in Table 1. However, there was a trend toward an interaction between genotype and sex identified by 2-way ANOVA for body fat percentage (genotype:sex p = 0.1166) where *Lgals3-*R200S^KIKI^ females had ∼4.3% higher body fat percentage and *Lgals3-*R200S^KIKI^ males had ∼5% lower body fat percentage than wild-type littermates (Table 1). Consistent with a reduction in galectin-3 secretion, we observed significantly reduced galectin-3 protein levels in the plasma of heterozygous and homozygous mutant mice (Table 1).

**Table 1.** Body composition and plasma galectin-3 levels of wild-type (*Lgals3*^+/+^), heterozygous (*Lgals3-*R200S^KI/+^), and mutant (*Lgals3*-R200S^KI/KI^) mice.^a^.

| Variables compared<sup>b</sup> | Females (36 wk) | Males (36 wk) |
| --- | --- | --- |
| **Body weight (g)** |  |  |
| Wild type | 31.26 ± 1.60 | 44.41 ± 1.64 |
| Heterozygotes | 34.15 ± 1.41 | 46.86 ± 1.74 |
| Mutant | 34.32 ± 1.22 | 44.13 ± 2.07 |
| **Body fat (%)** |  |  |
| Wild type | 32.19 ± 2.57 | 36.60 ± 1.94 |
| Heterozygotes | 36.26 ± 2.21 | 35.65 ± 1.51 |
| Mutant | 36.47 ± 2.40 | 31.57 ± 1.70 |
| **aBMD (g/cm<sup>2</sup>)** |  |  |
| Wild type | 0.057 ± 0.001 | 0.059 ± 0.001 |
| Heterozygotes | 0.057 ± 0.001 | 0.060 ± 0.001 |
| Mutant | 0.058 ± 0.001 | 0.059 ± 0.001 |
| **Plasma galectin-3 (ng/mL)** |  |  |
| Wild type | 29.7 ± 1.9 | 172.3 ± 22.4 |
| Heterozygotes | **22.2 ± 1.4\*\*** | **107.4 ± 22.4\*** |
| Mutant | **11.0 ± 1.4\*\*\*** | **29.8 ± 6.0\*\*\*** |

<sup>a</sup> Values are expressed as mean ± S.E.M. (n = 9-15). Dunnett's post-hoc analysis adjusted p-values compared to wild type; bold values highlight \*p < 0.05, \*\*p < 0.01, \*\*\*p < 0.001.
<sup>b</sup> aBMD = areal bone mineral density.

### Improved Trabecular Bone Parameters in Aged *Lgals3*-R200S Female Mice

In female mice with complete deletion of the *Lgals3* gene, we previously observed significantly increased cancellous bone mass in both the femurs and L3 vertebrae at 36 weeks [38]. As shown in Table 2, we observed a similar phenotype in mice where just the glycan binding domain of galectin-3 was disrupted. In L3 vertebrae, *Lgals3-*R200S^KIKI^ females had significantly elevated bone mineral density (BMD; +46.2%), bone volume fraction (BV/TV; +37.6%; Figure 2C), trabecular thickness (Tb.Th; +8.9%; Figure 2D), and trabecular number (Tb.N; +23.8%; Figure 2E) as well as decreased trabecular spacing (Tb.Sp; −9.7%; Figure 2F).

**Table 2.** Trabecular parameters of wild-type (*Lgals3*^+/+^), heterozygous (*Lgals3-*R200S^K*I*/+^), and mutant (*Lgals3-*R200S^KI/KI^) mice.^a^.

| Variables compared<sup>b</sup> | Females |  | Males |  |
| --- | --- | --- | --- | --- |
|  | Femur | L3 | Femur | L3 |
| **BMD (g/cm<sup>3</sup>)** |  |  |  |  |
| Wild type | 0.075 ± 0.010 | 0.133 ± 0.013 | 0.181 ± 0.009 | 0.213 ± 0.010 |
| Heterozygote | 0.065 ± 0.005 | 0.131 ± 0.007 | 0.206 ± 0.018 | 0.224 ± 0.017 |
| Mutant | 0.096 ± 0.012 | **0.186 ± 0.015\*** | 0.210 ± 0.016 | 0.219 ± 0.015 |
| **BV/TV (%)** |  |  |  |  |
| Wild type | 3.95 ± 0.58 | 11.72 ± 0.96 | 14.67 ± 0.82 | 19.37 ± 0.78 |
| Heterozygote | 3.07 ± 0.29 | 11.70 ± 0.66 | 16.59 ± 1.86 | 20.07 ± 1.62 |
| Mutant | 5.48 ± 0.91 | **16.08 ± 1.32\*** | 16.69 ± 1.29 | 16.08 ± 1.32 |
| **Tb.Th (mm)** |  |  |  |  |
| Wild type | 0.055 ± 0.002 | 0.056 ± 0.001 | 0.061 ± 0.002 | 0.055 ± 0.001 |
| Heterozygote | 0.055 ± 0.002 | 0.057 ± 0.001 | 0.062 ± 0.001 | 0.056 ± 0.001 |
| Mutant | 0.057 ± 0.002 | **0.061 ± 0.001\*** | 0.061 ± 0.002 | 0.053 ± 0.001 |
| **Tb.Sp (mm)** |  |  |  |  |
| Wild type | 0.408 ± 0.035 | 0.314 ± 0.011 | 0.246 ± 0.007 | 0.210 ± 0.005 |
| Heterozygote | 0.435 ± 0.019 | 0.310 ± 0.007 | 0.242 ± 0.014 | 0.214 ± 0.009 |
| Mutant | 0.383 ± 0.019 | **0.280 ± 0.011\*** | 0.233 ± 0.010 | 0.209 ± 0.008 |
| **Tb.N (mm<sup>-1</sup>)** |  |  |  |  |
| Wild type | 0.735 ± 0.110 | 2.083 ± 0.163 | 2.407 ± 0.116 | 3.508 ± 0.100 |
| Heterozygote | 0.558 ± 0.045 | 2.053 ± 0.086 | 2.627 ± 0.258 | 3.515 ± 0.236 |
| Mutant | 0.923 ± 0.131 | **2.613 ± 0.170\*** | 2.693 ± 0.177 | 3.677 ± 0.169 |

**Table 2 (Cont'd)**
| Variables compared<sup>b</sup> | Females |  | Males |  |
| --- | --- | --- | --- | --- |
|  | Femur | L3 | Femur | L3 |
| **Conn.D (mm<sup>-3</sup>)** |  |  |  |  |
| Wild type | 33.36 ± 5.73 | 64.55 ± 8.39 | 97.74 ± 7.08 | 120.25 ± 4.90 |
| Heterozygote | 22.99 ± 2.64 | 54.12 ± 3.15 | 110.63 ± 13.55 | 124.36 ± 12.38 |
| Mutant | 41.23 ± 7.82 | 87.38 ± 9.45 | **126.94 ± 13.06\*** | 139.49 ± 9.24 |
| **Bone length (mm)** |  |  |  |  |
| Wild type | 13.33 ± 0.16 | 3.60 ± 0.05 | 13.06 ± 0.06 | 3.67 ± 0.04 |
| Heterozygote | 13.22 ± 0.08 | 3.66 ± 0.04 | 13.22 ± 0.11 | 3.76 ± 0.05 |
| Mutant | 13.23 ± 0.09 | 3.67 ± 0.04 | 13.06 ± 0.11 | 3.74 ± 0.04 |

<sup>a</sup> Values are expressed as mean ± S.E.M. (n = 9-15); Holm-Sidak post-hoc analysis adjusted p-values compared to wild type; bold values highlight \*p < 0.05.
<sup>b</sup> BV/TV = bone volume/total volume; Tb.Th = trabecular thickness; Tb.Sp = trabecular spacing; Tb.N = trabecular number; Conn.D = connectivity density.

**Figure 2.**
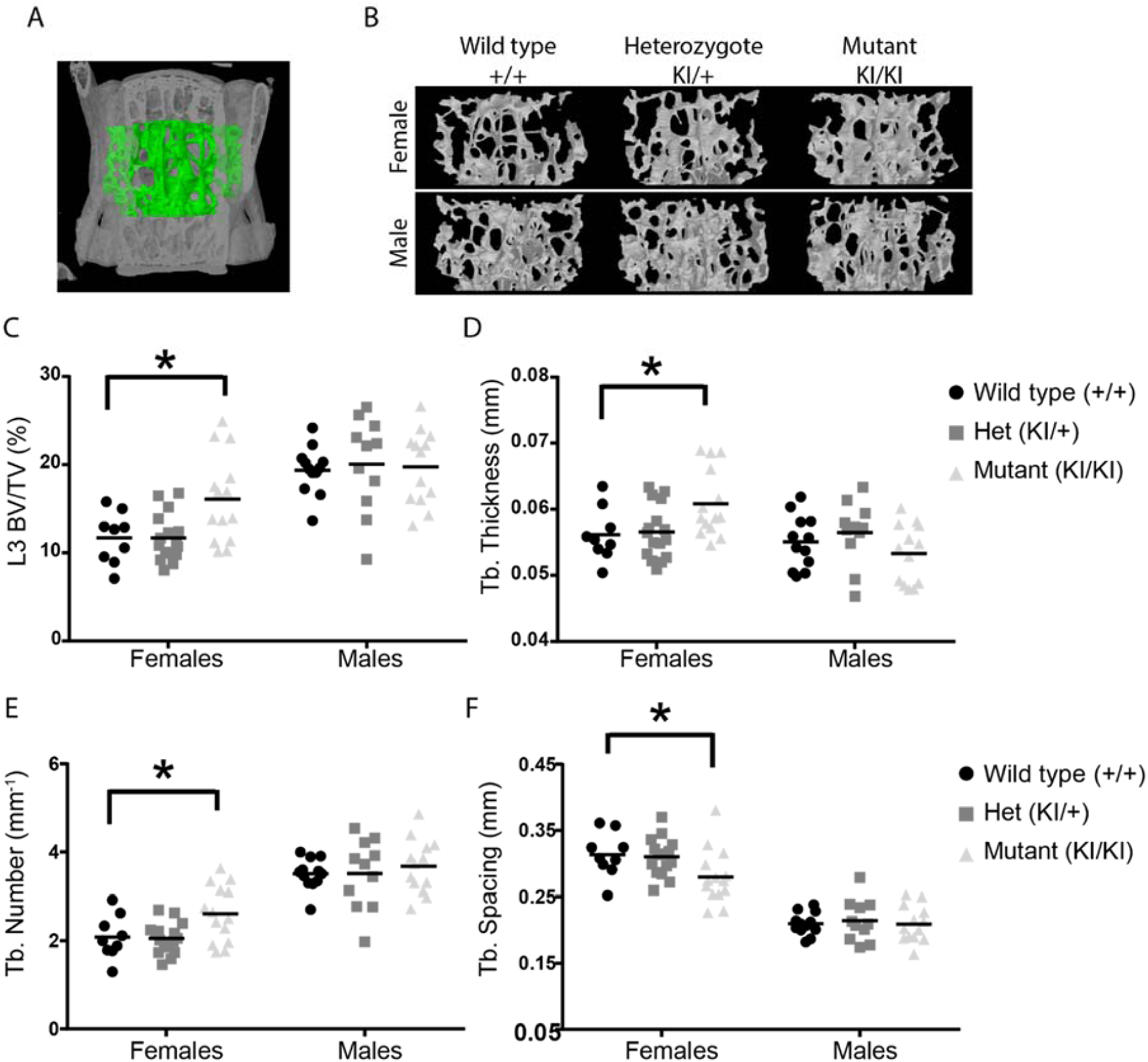
Enhanced trabecular bone mass at 36 weeks in *Lgals3-*R200S females. (A) Volume of interest for L3 trabecular bone measurements, as assessed by µCT. (B) Representative trabecular bone volumes of mice in each of the treatment groups. Scatter plots for bone volume fraction (BV/TV; C), trabecular (Tb.) thickness (D), Tb. Number (E), and Tb. Spacing (F). Asterisks indicate a significant difference, *p < 0.05.

Trabecular parameters in the distal femur trended in the same direction, but only *Lgals3-*R200S^KIKI^ males had a statistically significant effect with enhanced connectivity density (Conn.D; +29.9%; Table 2). The effect sizes observed in *Lgals3-*R200S^KIKI^ female L3 trabecular bone were similar to what we observed when we compared the L3 vertebrae of *Lgals3*^KOKO^ females to their wild-type littermates at this time point. However, the effect size of trabecular bone increase in the femurs of *Lgals3-*R200S^KIKI^ females compared to their wild-type littermates was lower than when we compared *Lgals3*^KOKO^ and wild-type femurs.

### Enhanced Cortical Bone Expansion in Aged *Lgals3*-R200S Male Mice

In both male and female mice with complete deletion of the *Lgals3* gene, we previously observed a significant increase in cortical bone expansion by 36 wk as reflected by increased total area (T.Ar) in male and female *Lgals3*-deficient mice, as well as significantly increased bone area (B.Ar) in female *Lgals3*-deficient mice [38]. We once again observed a similar cortical expansion phenotype in *Lgals3-*R200S mice, however only in males (Table 4 and Figure 3). *Lgals3-*R200S^KIKI^ males had a significant increase in T.Ar (+9.3%; Figure 3D) and marrow area (M.Ar; +11.9%; Figure 3F). *Lgals3-*R200S^KI+^ males had a significant increase in B.Ar (+9%; Figure 3E) compared to wild-type males. The effect size of the increased T.Ar and M.Ar in *Lgals3-*R200S^KIKI^ males was twice as strong compared to what we observed in *Lgals3*^KOKO^ male mice [38]. The effect size in *Lgals3-*R200S^KIKI^ females was roughly half of what we previously observed in *Lgals3*^KOKO^ females at this age[38].

**Table 4.**
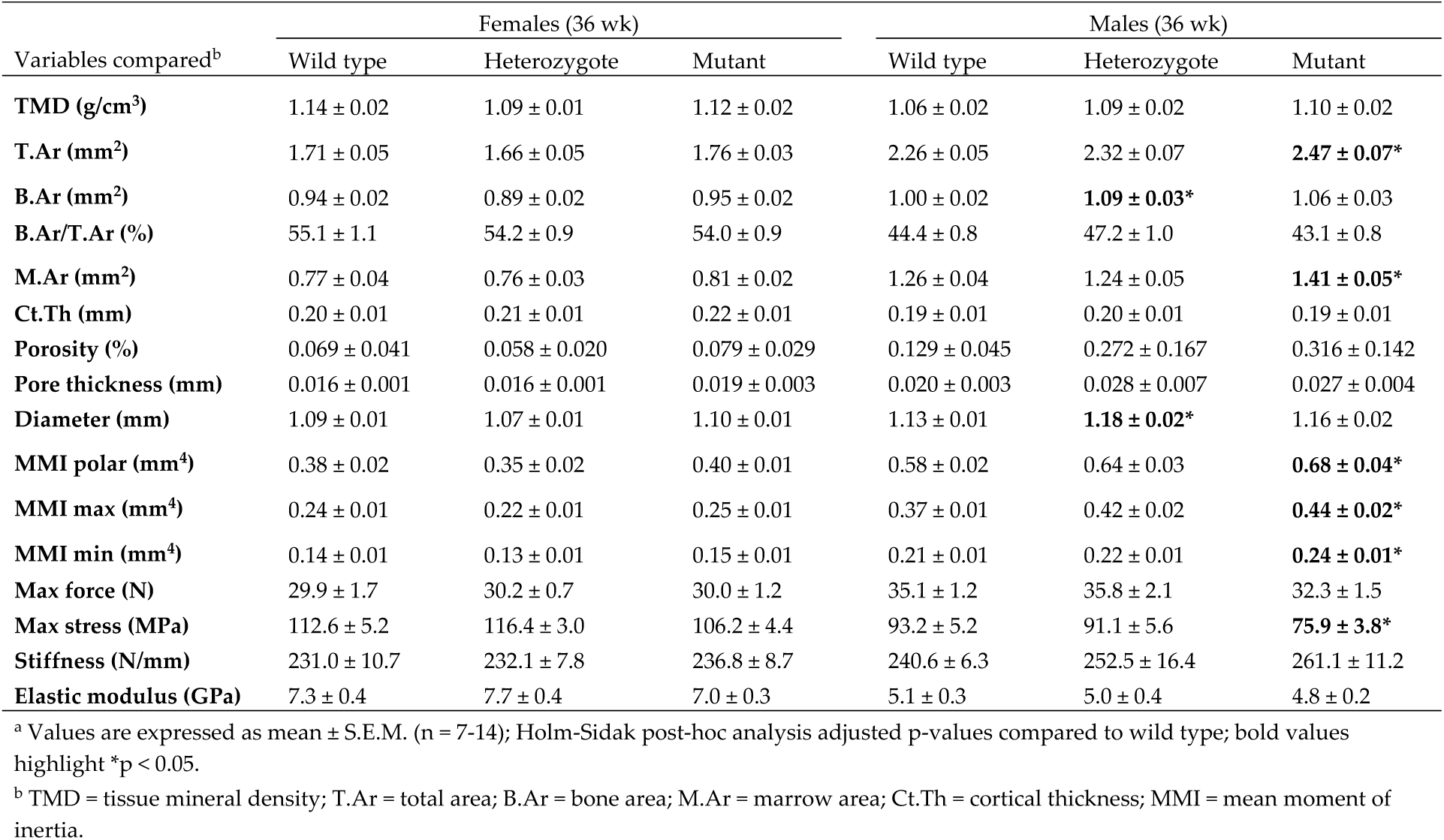
Femoral cortical bone parameters of wild-type (*Lgals3*^+/+^), heterozygous (*Lgals3-*R200S^KI/+^), and mutant (*Lgals3-*R200S^KI/KI^) mice.^a^.

**Figure 3.**
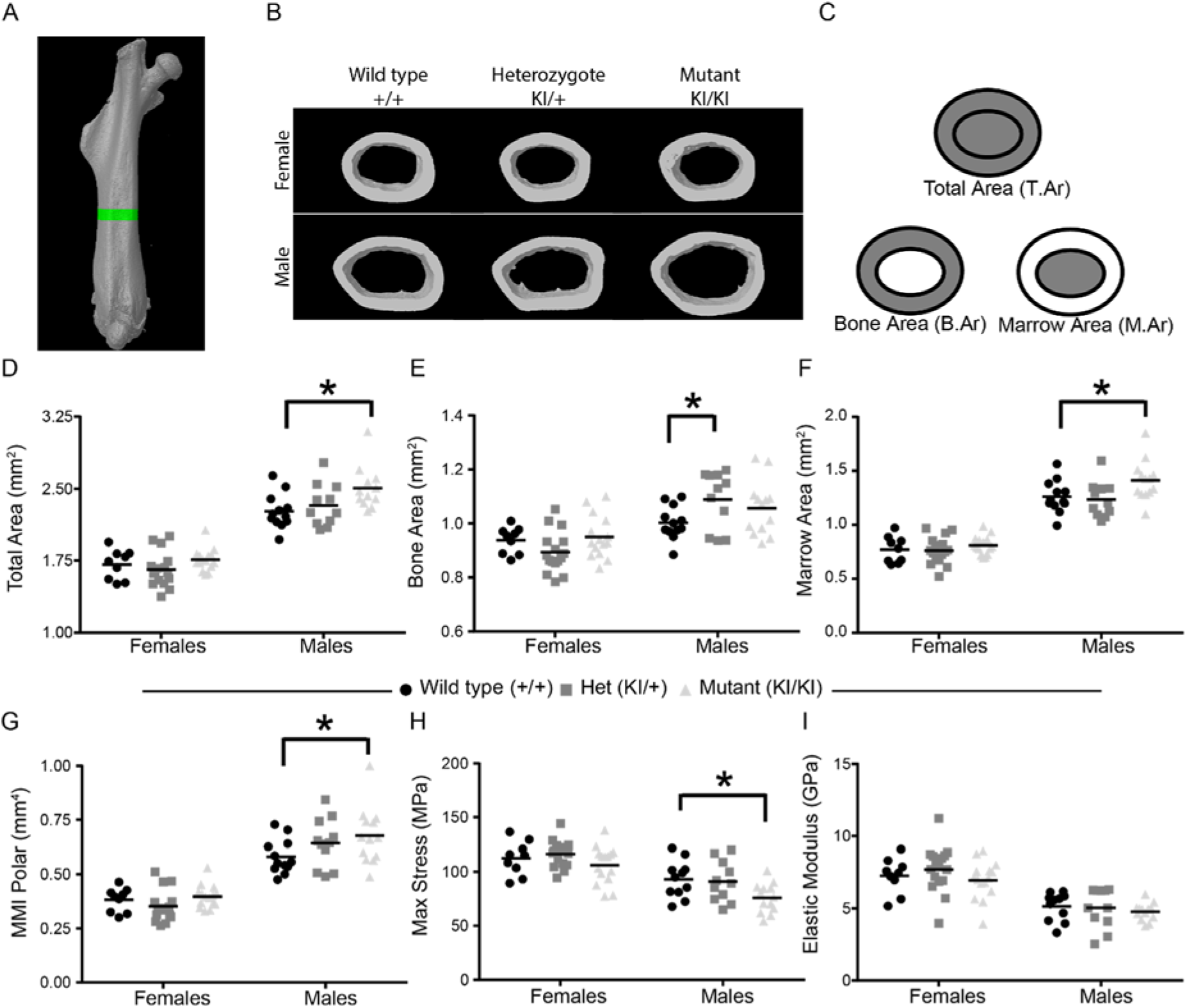
Enhanced cortical bone mass at 36 weeks in *Lgals3*-R200S males. (A) Region of interest for cortical bone measurements, as assessed by µCT. (B) Representative cortical bone volumes of mice in each of the treatment groups. (C) Diagram showing areas represented by total area, bone area, and marrow area. Scatter plots from µCT data for total area (D), bone area (E), marrow area (F), and mean polar moment of inertia (MMI Polar; G). Scatterplots from derived tissue mechanical properties measured by 4-point bending including max stress (H) and elastic modulus (I). Asterisks indicate a significant difference, *p < 0.05.

### Reduced Bone Quality in Aged *Lgals3*-R200S Male Mice

When we previously analyzed the mechanical properties of 36 wk *Lgals3*^KOKO^ female mice, we observed a non-significant decrease in tissue level strength (max stress) and a strong reduction in tissue level stiffness (elastic modulus). *Lgals3*^KOKO^ males trended the same direction, but the effect size was small [38]. This contrasts considerably with what we observed at this age in *Lgals3-*R200S^KIKI^ mice (Table 4 and Figure 3). Neither male nor female *Lgals3-*R200S mice showed a change in elastic modulus (Figure 3I) and trended toward increased whole bone stiffness (Table 4). Female *Lgals3-*R200S^KIKI^ mice also showed little-to-no reduction in max force (Table 4) or max stress (Figure 3H). Male *Lgals3-*R200S^KIKI^ mice, despite having significantly improved cortical geometry compared to wild-type males (MMI min: +14%, MMI max: +19%), had a trend toward reduced whole bone strength (Max force: −8%) and significantly reduced tissue level strength (Max stress: - 18.6%; Figure 3H).

## Discussion

In our previous studies we observed that mice with genomic loss of galectin-3 (*Lgals3*-KO mice), had increased cortical bone expansion between 24 and 36 weeks of age [38]. The effect size of the increases in cortical bone size were greater in *Lgals3*-KO females, as well as *Lgals3*-KO females had significant protection against age-related trabecular bone loss. The female dominance of the phenotype was further supported by use of a separate *Lgals3* null allele (*Lgals3*-Δ), where females once again had significantly increased trabecular and cortical bone mass at 36 weeks, but male *Lgals3*-Δ had slight reductions in both cortical and trabecular bone mass. The apparent sex-dependency of the bone phenotype was most likely due to diminished bone mass accrual in *Lgals3*-KO males prior to 12 wk of age[38, 39], which led us to suspect that *Lgals3*-KO mice might have reduced androgen-induced cortical bone expansion during puberty [40].

In this study we generated mice with a putative glycan binding deficient version of galectin-3 (*Lgals3*-R200S) in order to gain insight into whether the increased bone phenotype from *Lgals3*-KO mice was driven primarily by the loss of extracellular galectin-3. While we did not verify that our mutant lost glycan binding function, mutation of this arginine to a serine in human galectin-3 (R186S) has been shown to prevent galectin-3 secretion and glycan binding [30, 31, 37]. Because mutation of the functionally equivalent arginine in galectin-7 also prevents glycan binding [32, 33], we believe that the R200S mutation in galectin-3 should also be functionally equivalent. Future experiments will need to address whether this mutation in mouse galectin-3 is functionally equivalent to human galectin-3, as well as whether this mutation disrupts known intracellular interactions of galectin-3.

In *Lgals3*-R200S mice, we once again observed significantly increased trabecular bone mass in female, but not male mutant mice. This suggests that this phenotype in *Lgals3*-KO and *Lgals3*-Δ mice was primarily driven by loss of extracellular galectin-3. We did not perform histomorphometry or primary cell differentiation assays on these mice to confirm that this phenotype was driven primarily by an increase in osteogenesis as we did in *Lgals3*-KO mice.

While the trabecular phenotype was similar between *Lgals3*-KO/*Lgals3*-Δ and *Lgals3*-R200S mice, the cortical bone phenotype was a bit different. In *Lgals3*-KO mice we observed significantly increased T.Ar in both male and female mutant mice and increased B.Ar in female mice at 36 weeks. In *Lgals3*-Δ mice we similarly observed increased T.Ar and B.Ar in female, but not male mutant mice, supporting a female dominant phenotype in cortical bone mass [38]. However, in *Lgals3*-R200S mice, we observed a male dominant phenotype in cortical bone expansion. Our gonadectomy study suggested that global loss of galectin-3 may lead to reduced bioavailability of androgens[29]. This would reduce the ability of androgen to support bone mass accrual during puberty [40]. Altered sex-hormone regulation in the *Lgals3*-KO mother, during fetal development, might also explain why a different skeletal phenotype (increased age-related bone loss) has been reported when comparing *Lgals3*-KO mice to litters of background matched wildtype-mice [41]. More studies looking at systemic changes in hormones and growth factors in *Lgals3*-KO and *Lgals3*-R200S mice would help answer this question.

While we were unable to address more detailed mechanisms in this study, we believe that the *Lgals3*-R200S mice will help answer some of the important mechanistic debates about how galectin-3 influences signaling pathways and whether its role in these pathways is extracellular or intracellular. This will be accomplished by using cells from these mice that express physiologically relevant levels of galectin-3 that has low or no ability to function extracellularly. We look forward to sharing these mice with the galectin community in order to begin to unravel the complex functions of galectin-3.

## Acknowledgements

We thank other members of the Williams Lab for advice and assistance. Key members of the VARI Vivarium and Transgenics Core for this work include Bryn Eagleson, Audra Guikema, Tristan Kempston, and Tina Meringa. Support for this work was provided by the Van Andel Institute and the Van Andel Institute Graduate School. BOW receives support from a sponsored research agreement from Janssen Pharmaceutica and is a stockholder and member of the Scientific Advisory Board for Surrozen.

